# In vivo pulse-chase in *C. elegans* reveals intestinal histone turnover changes upon starvation

**DOI:** 10.1101/2025.02.13.638128

**Authors:** Christopher Borchers, Kara Osburn, Hyun Cheol Roh, Scott T. Aoki

## Abstract

The ability to study protein dynamics and function in the authentic context of a multicellular organism is paramount to better understand biological phenomena in animal health and disease. Pulse-chase of self-labeling fusion protein tags provide the opportunity to label proteins of interest and track those proteins over time. There are currently several challenges associated with performing in vivo protein pulse-chase in animals, such as cost, reproducibility, and accurate detection methods. The *C. elegans* model organism has attributes that alleviate many of these challenges. This work tests the feasibility of applying the Halo modified enzyme (HaloTag) for in vivo protein pulse-chase in *C. elegans*. HaloTag intestinal histone reporters were created in the worm and used to demonstrate that reporter protein could be efficiently pulse-labeled by soaking animals in ligand. Labeled protein stability could be monitored over time by fluorescent confocal microscopy. Further investigation revealed reporter protein stability was dependent on the animal’s nutritional state. Chromatin Immunoprecipitation and sequencing (ChIP-seq) of the reporters showed incorporation in chromatin with little change hours into starvation, implying a lack of chromatin regulation at the time point tested. Collectively, this work presents a straightforward method to label and track proteins of interest in *C. elegans* that can address a multitude of biological questions surrounding protein stability and dynamics in this animal model.

## Introduction

Life relies on a concerted orchestra of biological processes, with proteins as the primary workhorse. Proteins must perform specific functions and then be degraded when no longer required. Aberrant mutation or expression of proteins can have profound impacts on health and disease (1). Thus, developing methods to dissect protein dynamics are critical to understand the basic molecular mechanisms that govern life.

Protein pulse-chase is a versatile tool that can be used to study protein dynamics in cells and multicellular organisms. Original protein-pulse chase methods involved global, non-specific labeling of proteins using heavy or radioactive isotopes of amino acids (2–5). While this strategy can assess global protein stability, the spatial expression and dynamics across tissues is lost. The development of modified enzyme tags enable targeted protein in vivo pulse-chase (6–8). HaloTag is a modifed *R. rhodochrous* haloalkane dehalogenase with a mutation that traps the enzyme substrate as a covalent intermediate (9). Its fluorescent ligand signal is more stable and less sensitive to fixatives compared to conventional fluorescent tags (9, 10). Covalent bond formation allows saturated labeling and enzyme-ligand stability, enabling HaloTag and its ligands to function in pulse-chase methods. In this strategy, the “pulse” occurs during enzyme crosslinking to free ligand. Proteins can be “chased” by tracking the complex over time. HaloTag has reportedly higher fluorescent signal to other modified enzyme tags (10, 11), and there are many commercially available HaloTag ligands, allowing flexibility for different labeling strategies. Thus, HaloTag technology is highly amenable for targeted protein-pulse chase applications.

HaloTag-based in vivo pulse-chase methods have been previously used to study protein stability in cell culture (6, 7, 12, 13) and in mice (14, 15). While informative, cell culture studies lack the true context of authentic tissue in living organisms. Conducting HaloTag pulse-chase in organisms like mice involves both economic and technical challenges. The high cost and large quantities of Halo ligands required present significant financial barriers.

*C. elegans* is an established model organism with advantageous characteristics for in vivo pulse-chase methods. Site-specific gene editing methods are well established to easily add single-copy reporters or tag endogenous genes. The animal has a fast generational life cycle, well understood responses to different biological environments, and a transparent body for straightforward imaging (16). Some tissue in *C. elegans* are easily recognizable by light microscopy and thus do not need additional markers to identify them. For example, the intestine of *C. elegans* is a large, distinct, highly dynamic tissue that serves as an animal’s contact to the outside world. In addition to its primary role in metabolism, the intestine functions in immunity, protein secretion, and metal detoxification (17–21). At hatching, larvae have 20 intestinal cells (22) that undergo cell division and endoreduplication to produce 30-34 nuclei with a DNA content of 32C, or 16 times more than a normal diploid nucleus (23). Like all metazoans, this DNA is organized through chromatin organization by histones, nuclear proteins that compact DNA and play a central role in gene expression (24). The core repeating unit of chromatin is the nucleosome, an octamer of histone subtypes by which DNA is wound around (25). The *C. elegans* core histones share significant sequence identity to the human counterparts (26). Furthermore, many histone modifications and chromatin remodeling factors are conserved across metazoans, from worms to humans, implying organisms share significant chromatin biology (27). Metabolic changes have been associated with dynamic changes in chromatin structure (28), illustrating histones and chromatin play a critical role maintaining metabolic homeostasis.

This work explores the feasibility of performing in vivo pulse-chase in *C. elegans*. HaloTag intestinal histone protein reporters were designed and could be labeled in vivo for imaging-based monitoring. Stability of labeled histone proteins changed with nutritional stress. The histone reporter proteins incorporated into chromatin, implying use in gene regulation. Collectively, this work illustrates the potential of *C. elegans* pulse-chase to study protein dynamics and function in animal biology and pathology.

## Results

### In vivo pulse labeling of intestinal histone reporters

The overarching goal was to develop a strategy to track the tissue specific stability of a target protein to determine its lifespan in vivo. Histone proteins were selected due to the extensive research of their assembly and function, and the *C. elegans* intestine selected due to its strikingly large nuclei and high DNA content in adult worms (23) (**Fig 1A**). Intestinal histone reporters were created using MosSCI gene editing (29) to insert a single copy of a reporter gene (**Fig 1B**, see **Methods**). The reporters consisted of an intestinal *elt-2* promoter (30); *his-58*, *his-9*, or *his-72*; a V5 epitope (31) and HaloTag (9), and an *unc-54* 3’UTR (**Fig 1B**). The three histone variants were chosen due to their unique functions and conservation across metazoans. *his-58* is a canonical H2B protein and a core nucleosome component (24). *his-9* and *his-72* are histone H3 variants that are commonly associated with gene repression and expression, respectively (24). The transgenes enabled tissue specific expression with identical gene regulatory elements. Thus, all protein expression level differences observed between histones would be due to inherent properties of the protein, not other factors associated with post-transcriptional regulation. Histone reporter worms fixed and stained with Halo-ligand showed specific expression in intestinal nuclei (**Fig 1C**), as expected of histones. Of note, *his-58*/H2B and *his-9*/H3.1 reporters showed robust fluorescence (**Fig 1C**), while *his-72*/H3.3 had fainter signal (**Fig S1A**). This difference in histone protein expression was also observed by immunoblot (**Fig S1B-C**). *his-9*/H3.1 and *his-72*/H3.3 differ by only nine amino acids (**Fig S1D**), but the proteins had significantly different protein expression levels. Due to the equivalent protein expression and research goals, the remainder of the study focused on using the *his-58*/H2B and *his-9*/H3.1 reporters to test protein labeling and tracking in vivo.

**Figure 1.**
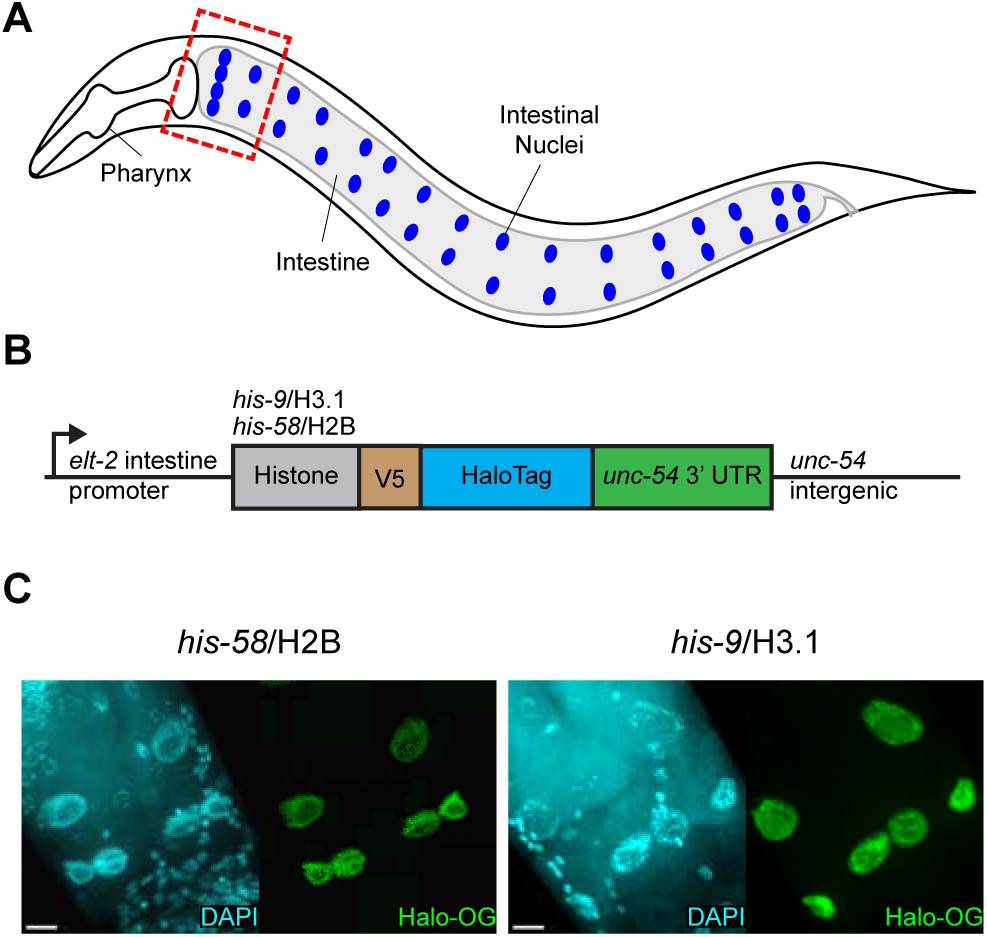
Intestinal histone reporters are robustly expressed. (**A**) Diagram of adult *C. elegans* intestinal nuclei. Animals have 20 intestinal cells and 30-34 nuclei (blue, 34 shown). Dashed box outlines intestinal region of interest in this study. (**B**) Linear diagram of the HaloTag intestinal histone reporter. An *elt-2* intestine-specific promoter drives histone variants fused to a V5 epitope and HaloTag in C. elegans. All reporters also have an *unc-54* 3’UTR and intergenic region. (**C**) Confocal images of histone reporter variant strains. Worms were stained with DAPI to visualize nuclei and Halo-Oregon Green (Halo-OG) visualize reporter protein. Images taken of the anterior portion of the intestine at 63X magnification under oil immersion and shown as max intensity projection of confocal stack. Scale bar, 10 µm.

The HaloTag enzyme variant covalently binds to haloalkane ligands (9). As demonstrated in (6, 7, 12, 13) and in vivo in multicellular organisms (14, 15), fluorescent ligands can be pulsed to label tagged proteins. To test this system in *C. elegans*, *his-58*/H2B reporter worms were soaked in different fluorescent HaloTag ligands, washed, fixed, counterstained with a different HaloTag ligand, and imaged by confocal microscopy (**Fig S2A**). Worms pulse-labeled with the Halo-TMR ligand had significantly higher fluorescence compared to the other ligands tested (**Fig S2A-B**). Pulse-labeled reporter worms were next evaluated for their efficiency in labeling nuclei throughout the intestine. *his-58*/H2B reporter worms were soaked with Halo-TMR, washed, fixed, counterstained, and the whole worm imaged by confocal microscopy (**Fig S3A**). While the intestinal histone reporter was uniformly expressed throughout the intestine (**Fig S3A-B**), Halo-TMR fluorescence signal was highest in nuclei at the anterior and posterior regions (**Fig S3A-B**). *C. elegans* have 30-34 intestinal nuclei, with variation dependent on variable division of the posterior nuclei (22, 23). Subsequent pulse-chase imaging experiments focused on the anterior intestinal nuclei to avoid this variability. In sum, this pulse-labeling strategy with Halo-TMR ligand enabled tissue-specific protein labeling. The strong fluorescent signal created the opportunity to test whether the labeling method could be used to follow protein stability over time.

To assess the feasibility of in vivo pulse-chase with the HaloTag in *C. elegans*, young adult *his-58*/H2B and *his-9*/H3.1 reporter worms were pulse-incubated with Halo-TMR ligand and then allowed to propagate on bacterial plates for several hours before collection, fixing, staining, and imaging (**Fig 2A**, see **Methods**). Worms that were either not pulse-labeled or collected immediately after labeling served as negative and positive controls, respectively. As observed previously, both *his-58*/H2B and *his-9*/H3.1 reporter protein were labeled with Halo-TMR ligand in worms at appreciable levels compared to controls (**Fig 2B, C**; **Fig S4A, B**). By 3 hours, Halo-TMR signal was significantly lost, reaching basal levels (**Fig 2B, C**; **Fig S4A, B, D**). Counterstaining of the reporters with a separate fluorescent ligand confirmed that the reporter was expressed in all worm samples (**Fig 2B; Fig S4A, D**). These findings demonstrate that the propagation of pulse-labeled animals lead to the loss of the pulse-labeled protein signal in *C. elegans*.

**Figure 2.**
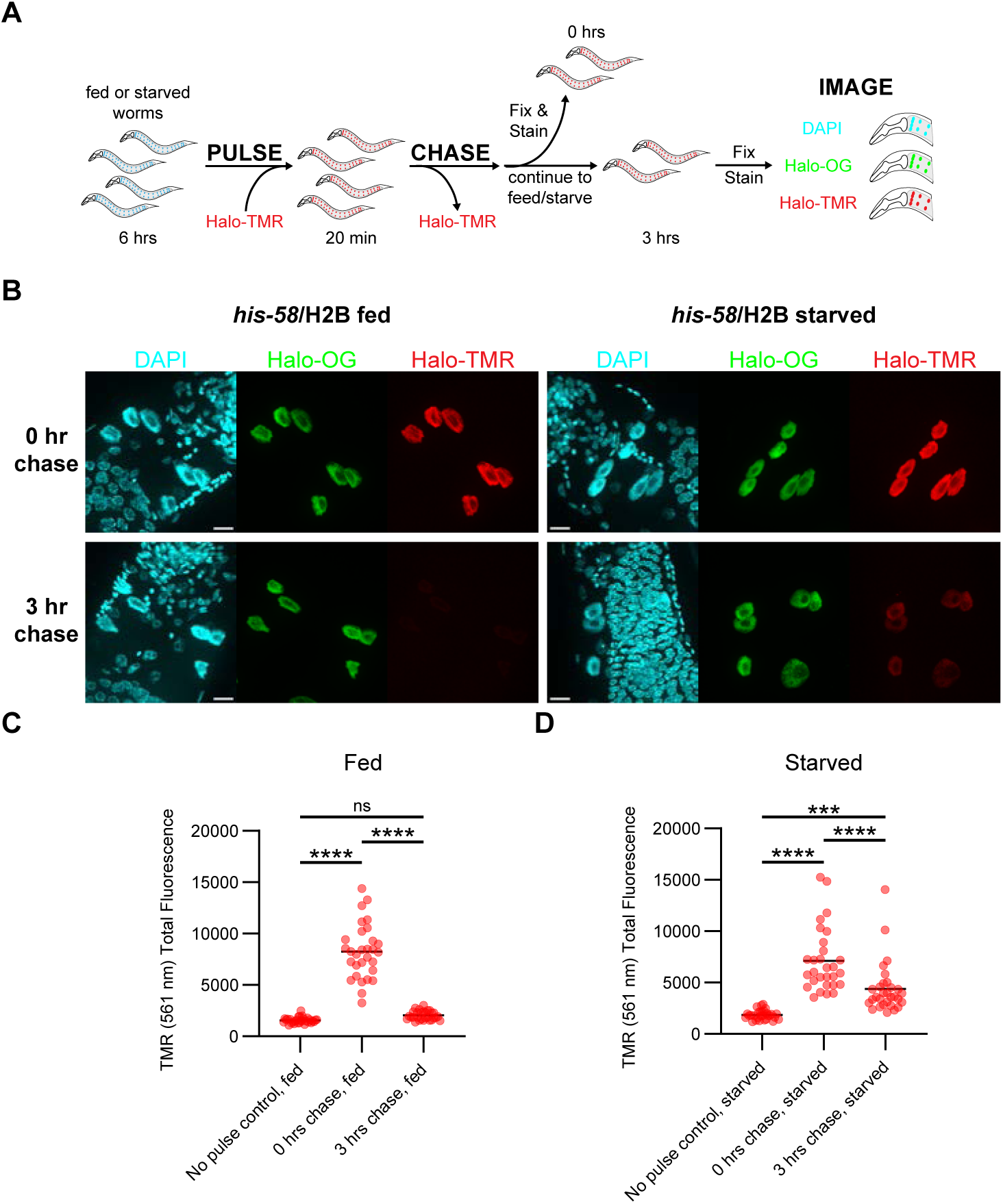
In vivo labeling of intestinal histone reporters. **(A)** In vivo HaloTag pulse-chase method. Intestinal histone reporter animals were pulse-labeled by soaking in the presence of Halo-TMR ligand. Animals were washed and allowed to propagate for 0 or 3 hours. At each timepoint, animals were collected, fixed, stained with DAPI (DNA) and Halo-Oregon Green (Halo-OG), and imaged using confocal microscopy to assess fluorescence intensity over time. (**B**) Representative images of fed and starved his-58/H2B reporter anterior intestinal nuclei after a 0- or 3-hour chase. Images shown as max intensity projection of confocal stack taken at 63X magnification under oil immersion. Scale bar, 10 µM. (**C-D**) Fluorescence intensity measurements of fed (C) and starved (D) his-58/H2B reporter anterior nuclei. Fluorescence was significantly higher after a 3-hour chase when compared to a no pulse control. Significance was determined by performing a two-way ANOVA with Tukey’s multiple comparisons test; ***p<0.001; ****p<0.0001; ns, not significant.

A concern was that the loss of signal observed was solely due to photobleaching or other confounders not connected to biology. As a control, an additional HaloTag nuclear protein in the intestine was pursued. Nuclear lamins are intermediate filament proteins that assemble the nuclear lamina, a structure that connects the nuclear envelope to chromatin (32). Lamins help sequester heterochromatin to the nuclear periphery and function in gene regulation (32). The proteins are stable in the intestine tissue, with an estimated half-life of 5 days in mice (33). *C. elegans* have one lamin gene, *lmn-1* (34). An LMN-1 intestinal reporter was generated by MosSCI single gene editing (29) (**Fig S5A**). The reporter design was similar to the other intestinal reporters, with an *elt-2* promoter (30), *lmn-1* exons and introns, V5 epitope (31), HaloTag (9), and *unc-54* 3’UTR and intergenic region (**Fig S5A**). Intestinal *his-58/*H2B and LMN-1 reporters were pulse-labeled by soaking worms in Halo-TMR, followed by washing and immediate collection or collection after 3-hours on bacteria-containing plates (**Fig S5B**). Worms were fixed, counterstained, and imaged by confocal microscopy. Fluorescence quantitation revealed that the mean fluorescence intensity was similar between *his-58*/H2B and LMN-1 reporters immediately after pulse-labeling (**Fig S5C, D**). However, after a 3-hour chase, the fluorescence intensity of the LMN-1 reporter was higher compared to the *his-58*/H2B reporter (**Fig S5C, D**). This result suggested that the LMN-1 reporter protein was more stable than histone H2B and provided supportive evidence that Halo tag pulse-chase method detected different protein stability with different proteins in vivo.

Previous studies demonstrated that starving worms affected chromatin in non-intestinal cell types (35). To probe the pulse-chase method further, reporter worms were starved to examine how nutritional stress influenced labeled protein stability. Young adult *his-58*/H2B and *his-9*/H3.1 reporter worms were starved prior to in vivo pulse-chase labeling and chasing in the absence of bacterial food (**Fig 2A**, see **Methods**). As performed previously, non-pulsed worms and worms collected immediately after pulse-labeling served as negative and positive controls, respectively. Starved reporter worms expressed the HaloTag histone reporters at comparable levels as fed worms, measured by immunoblot (**Fig S1B, C**), and were effectively pulse-labeled, determined by imaging (**Fig 2B, D; Fig S4A, C**). However, instead of returning to basal levels as seen in fed animals (**Fig 2B, C; Fig S4A, B, D)**, starved animals retained appreciable levels of pulse-labeled histone reporter protein signal (**Fig 2B, D; Fig S4A, C, D**). Collectively, the results suggested that the stability of reporter histone proteins were influenced by the animal’s nutritional status.

### Histone reporter proteins incorporate into chromatin

A concern using the histone reporters was their utility in the intestine. While the reporter histones colocalized with DNA in intestinal nuclei (**Fig 1**), it was unclear if the proteins were deposited into chromatin for use in gene regulation. To probe functionality, Chromatin Immunoprecipitation and Sequencing (ChIP-seq) was performed on fed or starved adult worms (**Fig. 3A**). After a 6-hour period of feeding or starvation, timing that aligned to pulse-chase experiments, adult reporter worms were collected and frozen. These worm samples were homogenized, crosslinked, sonicated to shear DNA, and reporter histone protein immunoprecipitated (see **Methods**, **Fig S6A**). As a negative control, the protocol was also performed on N2 wild type worms. Immunoprecipitated DNA was detected in all V5 test ChIP samples but not in the N2 control (**Table S1A**). Input and immunoprecipitated DNA were sequenced to identify genomic regions bound by the histone reporters (see **Methods**). Histone reporter peaks were identified in test samples versus input and N2 control background (**Fig 3B**) and were consistent with characteristics typically observed in histone ChIP-seq (36, 37). Additionally, multidimensional scaling (MDS) analysis of the identified peaks revealed unique chromatin binding profiles for each histone reporter (**Fig 3C**). Thus, these analyses were consistent with functional histone reporter proteins forming specific chromatin interactions in the intestine.

**Figure 3.**
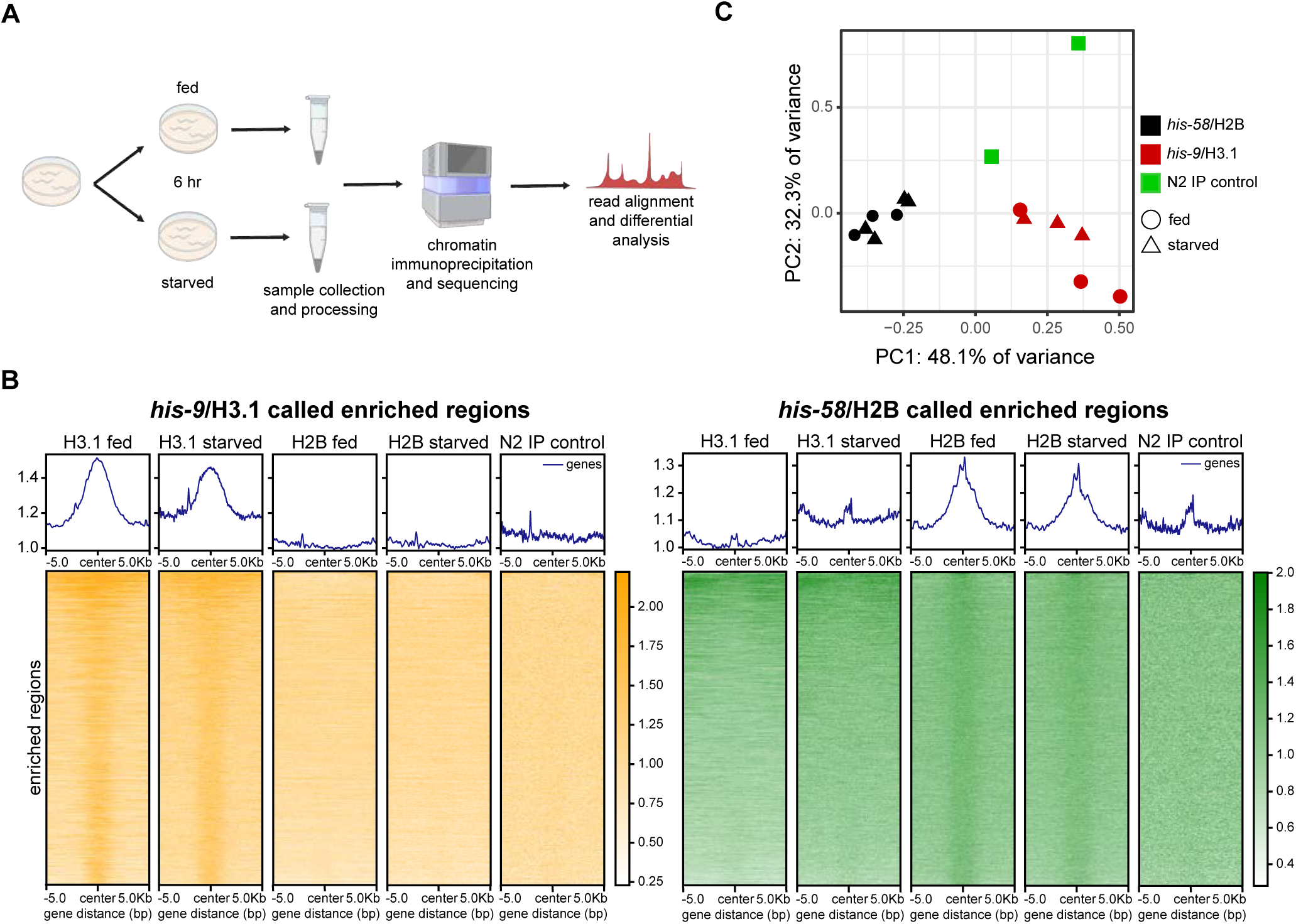
Intestinal histone reporters incorporate into chromatin. (**A**) ChIP-seq experimental workflow. Synchronized young adult worms were fed or starved for 6 hours prior to sample processing and sequencing. (**B**) Heatmap of identified *his-9*/H3.1 (left) and his-58/H2B (right) enriched regions identified by epic2. Enriched regions for his-9/H3.1 are absent in *his-58*/H2B and N2 control. Similarly, enriched regions in *his-58*/H2B were absent in *his-9*/H3.1 and N2 control. (**C**) Multidimensional Scaling (MDS) plot comparing identified enriched regions in *his-9*/H3.1, *his-58*/H2B, and N2 control samples. Samples cluster into respective strain groups. MDS analysis performed using edgeR program.

The broad nature of the identified histone peaks made specific gene target identification challenging. Instead, gene targets of the histone reporters were identified by a window-based strategy of signal enrichment within protein coding gene regions previously described for ChIP-seq analysis in other studies (38, 39). Using this strategy, analysis of *his-58*/H2B samples compared to negative controls revealed no significant gene targets, indicating evenly distributed peaks across the genome without specific regional enrichment. When *his-9*/H3.1 samples were compared to negative controls, the analysis identified 187 and 25 genes with significant histone reporter binding in fed and starved samples, respectively (**Fig 4A, B; Table S2**). All 25 starved targets were present in fed samples (**Fig 4A; Table S2**). When *his-9*/H3.1 fed and starved samples were compared, no genes with significant changes were identified. Thus, the *his-9*/H3.1 gene targets identified were only specific when compared to the controls, implying no major chromatin changes between animal nutritional states. Tissue Enrichment Analysis (TEA) identified that these *his-9*/H3.1 targets enriched for the intestine and other alimentary system related tissues (See **Methods**, **Fig 4C**), while Gene Ontology (GO) enriched for immune and metabolic pathways (See **Methods**, **Fig 4D**), associated functions of the *C. elegans* intestine (40). In sum, ChIP-seq revealed chromatin incorporation of histone reporters. The similarity of *his-9*/H3.1 gene targets between fed and starved animals suggested minimal chromatin changes at this time point of animal starvation.

**Figure 4.**
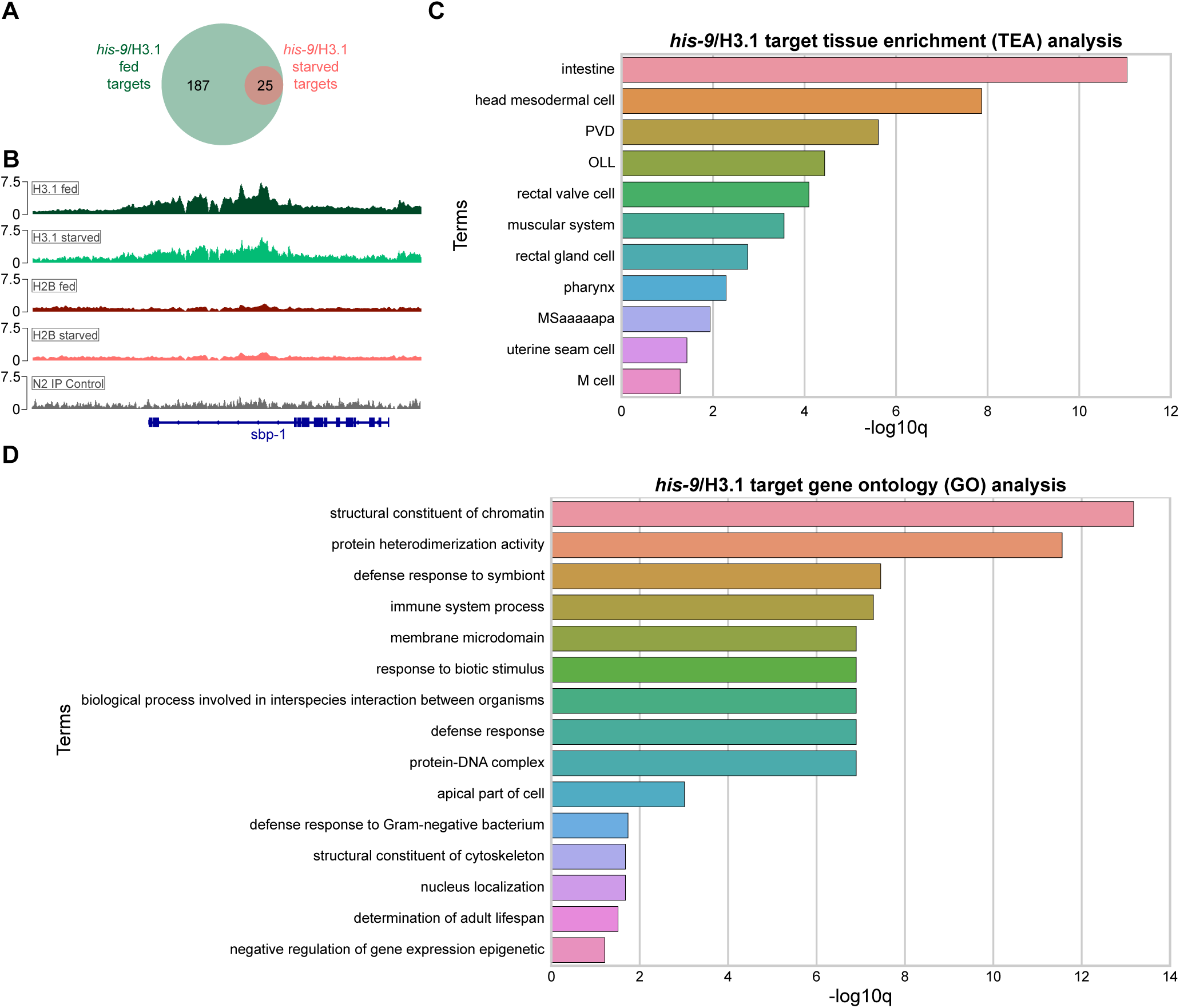
*his-9*/H3.1 identified targets enrich for intestine and intestine-related processes. (**A**) 187 genes were identified as *his-9*/H3.1 targets in fed animals and 25 genes were identified as *his-9*/H3.1 targets in starved animals. All 25 starved targets were present in fed samples. (**B**) Integrative Genomics Viewer (IGV) (76) browser view of *his-9*/H3.1 identified target, *sbp-1*. Samples were normalized to themselves using the Reads Per Genomic Content (RPGC) normalization method. Representative *his-9*/H3.1 fed and starved (green), *his-58*/H2B fed and starved (red), and N2 control (grey) sample shown. (**C**) Tissue enrichment analysis (TEA) of *his-9*/H3.1 identified targets. Alimentary system tissues enriched in both fed and starved animals. (**D**) Gene Ontology analysis (GO) of *his-9*/H3.1 identified targets. Terms in both fed and starved samples reflect alimentary system-related processes.

## Discussion

Monitoring the stability and localization of proteins in animals necessitates the ability to label and track a specific protein population across space and time. This work used the HaloTag modified enzyme to establish an *in vivo* protein pulse-chase method in *C. elegans*. HaloTag reporter proteins in the intestine were efficiently labeled by exposing worms to a fluorescent HaloTag ligand. The stability of these pulse-labeled proteins could be tracked over time in animals, and protein stability was dependent on the animals’ nutritional state. Given that the reporter levels were similar in both fed and starved animal conditions, the increase in protein stability also implied a decrease in new protein synthesis. These histone reporter proteins were functional in that they incorporated into chromatin and bound broadly across genes, as expected of histone proteins. While no targets were able to be identified for histone *his-58*/H2B, histone his-9/H3.1 enriched for genes associated with intestine-associated processes. Thus, this *C. elegans* in vivo pulse-chase method enabled tracking of biologically-relevant proteins in specific animal tissue.

HaloTag-based, in vivo protein pulse-chase methods have been previously used in cell culture cell culture (6, 7, 12, 13) and in mice (14, 15). While informative, cell culture studies lack the context of authentic tissue, and working in mice elicits both economic and technical challenges. Ensuring reproducible ligand delivery can be difficult and may confound experimental results. This work demonstrated that HaloTag in vivo pulse-chase experiments were both straightforward and cost-effective in *C. elegans*. Delivery of the ligand was achieved by soaking the worms in membrane-permeable ligand. The transparent worm also made detection by imaging easy. Limitations also exist. In *C. elegans*, and multicellular animals generally, the current method can only measure protein dynamics on a timescale of hours to days. Studying the stability and localization of highly dynamic proteins remains a challenge *in vivo*. Efforts to mitigate this limitation include the development of non-fluorescent HaloTag ligands, which can be used in a dual “pulse” strategy to improve labeling timepoints (12, 41). Challenges of preferential uptake and low membrane permeability for certain tissue (such as the brain) have been reported in mice (15, 42). This work focused on labeling of HaloTag fusion proteins exclusively in the intestine of adult animals. Successful delivery and uptake of ligands by other tissues and during other developmental stages of the worm may be a limitation, which could directly affect labeling efficiency and utility of the method. HaloTag is also a relatively large tag (∼33 kDa) and thus requires functional characterization to ensure that the tag does not interfere with protein function. Regardless, the method holds promise to investigate longer lived proteins (43–48) and those that traffic across cells and tissue in animal development (49–51).

Histone H3.1 is well characterized and is reported to be deposited into chromatin in a replication-dependent manner (24, 52). The adult *C. elegans* intestine is a non-dividing tissue. The organ is fully formed by later developmental stages and serves the animal throughout life (17, 22). Only moderate H3.1 chromatin remodeling may occur during adulthood, which could partially explain the absence of dramatic H3.1 chromatin changes in starved adult animals. Alternatively, chromatin remodeling may robustly occur, however longer starvation periods may be necessary to detect changes in *his-9*/H3.1 chromatin targets. Changes to chromatin architecture in response to starvation are reported in *C. elegans* (35, 53) and other organisms (54). Transcriptomic analyses of *C. elegans* starved for durations similar to this study showed only modest changes compared to those starved for significantly longer periods (40). Furthermore, worms starved for longer periods had chromatin architecture changes in other cell types (35). Future work will investigate the sites of histone chromatin assembly across different types and durations of animal stress states.

Developing methods to study protein dynamics *in vivo* is critical to understanding biological processes *in situ*, or their authentic context. This work serves as entrée to illustrate the potential of in vivo pulse-chase methodologies in *C. elegans.* From pulse-labeling to detection, the method is highly adaptable to cater investigation into a wide variety of biological processes *in vivo*. For example, proteome dysfunction and aggregation are hallmarks of aging and neurological disorders (reviewed in (55, 56)). HaloTag pulse-chase could be used to directly observe and measure protein stability and aggregation in the worm. The method could also be applied toward understanding how RNA-protein inheritance dictates transgenerational epigenetic inheritance (reviewed in (57)). Addition of other modified enzyme tags, like SNAP (58), will make it possible to label and track two different proteins of interest simultaneously, further expanding the experimental application. Technical advancements made in *C. elegans* will serve as a stepping-stone to address a variety of questions in metazoan biology.

## Materials and Methods

### Worm maintenance

*C. elegans* were grown and maintained on nematode growth medium (NGM; 25 mM KPO4 pH 6.0, 5 mM NaCl, 1 mM CaCl_2_, 1 mM MgSO_4_, 2.5 mg/ml tryptone, 5 µg/ml cholesterol, 1.7% agar) seeded with *E. coli* HB101 as previously reported for standard nematode growth conditions (59). To isolate genomic DNA for genotyping, worms were picked into worm lysis buffer (10 mM Tris-HCl pH 8.3, 50 mM KCl, 2.5 mM MgCl_2_, 0.45% Triton X-100, 0.45% Tween-20, 0.01% Gelatin) containing Proteinase K (200 ug/mL) and lysed at 60°C for 1 hour.

### Reporter generation

The *elt-2* promoter and genes of interest (*his-9*, *his-58*, *his-72*, *lmn-1*) were Q5 (NEB, #M0491S) PCR amplified from worm lysate. A *C. elegans* optimized HaloTag (9) was generated by commercial gene fragment synthesis (Twist Biosciences), and the V5 tag (31) was included in PCR primers. Q5 PCR amplified fragments were ligated into pCFJ151 (29) MosSCI vector using Gibson cloning (60). Injections were performed using an Eclipse Ts2 microscope (Nikon), IM-300 Microinjector (Narishige), and Microscope Mechanical Micromanipulator (Leica). EG6699 (*ttTi5605* II) (61) uncs were injected with pDDC14, pDDC15, pDDC16, or pDDC171, somatic RFP markers (pGH8, pCFJ90, pCFJ104 (29)) and the germline transposase, pCFJ601 (61) following MosSCI direct insertion protocol (29). The transgenic animals with Mos site insertions rescued the *unc* phenotype (*unc-119*) and lost the somatic RFP reporters. Transgene insertion was validated by PCR and proper repair determined by DNA sequencing. All generated strains were outcrossed twice with N2 worms. See **Table S3** for plasmids and strains used in this study.

### Pulse Labeling

Synchronized reporter worms were washed in Phosphate Buffer Saline (PBS) + 0.01% Tween-20 (PBS-T) to remove excess bacteria and soaked for 20 minutes in the presence of 300 nM Halo-TMR (G8251, Promega; Madison, WI), Halo-R110Direct (G3221, Promega, G3221; Madison, WI), or Halo-Oregon Green (G2801 Promega; Madison, WI). Worms incubated without HaloTag ligand served as a negative control. Worms were washed with PBS-T to remove excess ligand. The worms were then further propagated on NGM plates without or with HB101 bacteria, or immediately fixed in 3% Paraformaldehyde (PFA; 50-980-487, Fisher) and 100% ice cold Methanol (-20°C). Propagated worms were later collected at specific timepoints, washed with PBS-T, and fixed as described above. All worms were stained with 1:2000 DAPI to visualize nuclei and counterstained with a different HaloTag ligand to visualize total reporter protein. Worms pulse-labeled with Halo-TMR were counterstained with Halo-Oregon Green and worms pulse-labeled with either Halo-R110Direct or Halo-Oregon Green were counterstained with Halo-TMR. Samples were pipetted onto slides in Vectashield (H-1900-10, Vector Laboratories Inc; Newark, CA), covered with a glass coverslip, and sealed with nail polish. Slides were stored at 4°C until imaging. Worms were imaged by spinning disk confocal microscopy on a Zeiss Axio Observer Z1 modified by 3i (www.intelligent-imaging.com).

### Image Processing/Quantification

Raw image files were loaded into ImageJ FIJI (62) and fluorescence levels normalized between different ligand groups. To quantify fluorescence, a mask was generated from the Halo counterstain channel image to exclusively mark the intestinal nuclei. The fluorescent threshold levels used for mask generation was maintained between sample groups. The mask was used to quantify the mean fluorescence value of the pulse-labeled Halo ligand channel for the first six nuclei of the intestine. Oversaturated images or poorly outlined nuclei were not used in fluorescence quantitation. Fluorescence quantitation for *lmn-1* reporter animals (*ddcSi43 [elt-2 promoter::lmn-1::histone linker::1xV5::Halo opt with introns::unc-54 3’UTR and intergenic region] II*) followed the same protocol as histone reporters. A mask was generated from the Halo counterstain channel image to mark *lmn-1* puncta in intestinal nuclei. The mask was used to quantify the mean fluorescence of the pulse-labeled *lmn-1* Halo ligand channel in the first six nuclei of the intestine. Mean fluorescence of *lmn-1* was then compared to the mean fluorescence of CDE6 (*ddcSi2 [elt-2 promoter::his-58::histone linker::1xV5::Halo opt with introns::unc-54 3’UTR and intergenic region] II*). Comparison of the fluorescence levels of the different ligands was performed in GraphPad Prism 10.4.1 (GraphPad Software, Inc., San Diego, CA).

### Immunoblots

Histone reporter strains were grown on NGM plates seeded with HB101 bacteria. Plates from each strain were bleached to synchronize strains at the L1 larval stage (63). L1 larvae were plated NGM plates with HB101 and grown for 72 hours (24 hours-post L4 stage). Worms were washed from the plate with PBS-T, pipetted into conical tubes, and washed with PBS-T to remove remaining bacteria. After the final wash, the samples were pipetted onto NGM plates without or with HB101. These samples were incubated for 6 hours at 20°C and again washed with PBS-T to remove bacteria. Samples were transferred to tubes with 5x SDS-PAGE sample buffer (250 mM Tris pH 6.8, 25 mM EDTA pH 8.0, 25% glycerol, 5% SDS), frozen at -80°C, and stored at -20°C until use. Samples were run on SDS-PAGE gel and transferred to PVDF (BioRad, #1620264) using the BioRad TransBlot Turbo System (BioRad, #1704150). Blots were probed with a primary anti-V5 monoclonal antibody (R&D Systems Bio-techne-NBP1-80562) and secondary goat-anti-mouse Horse Radish Peroxidase antibody (VWR, #95058-740) prior to incubation in SuperSignal Plus West Femto (Thermo, #34094) and SuperSignal Plus West Pico (Thermo, #34580). Blots were imaged on a BioRad ChemiDoc MP system (BioRad, #12003154). Blots were stripped and re-probed with a primary anti-GAPDH antibody (60004-1-Ig) and secondary goat-anti-mouse Horse Radish Peroxidase antibody (VWR, #95058-740). Chemiluminescence exposure settings were kept identical between replicate blots to maintain reproducibility.

Quantification of protein bands was performed using Image Lab (BioRad) and ImageJ FIJI (62). In Image Lab, chemiluminescent image contrast was adjusted to optimal, identical levels. ImageJ FIJI was used to determine band density. Relative protein expression of the histone reporters were determined by a quotient of histone reporter over GAPDH loading control band density. Statistical significance was determined using a two-way analysis of variance (ANOVA) and Tukey’s multiple comparisons test in GraphPad Prism 10.4.1.

## Chromatin ImmunoPrecipitation and sequencing (ChIP-seq)

### Sample Collection

CDE5, CDE6, and N2 worms were bleached for a synchronous population of L1 larvae (63). L1s were pipetted onto NGM plates with concentrated HB101 and propagated for 72 hours. Worms were collected and washed with PBS-T to remove bacteria then added to concentrated HB101 or empty plates. These worms were fed or starved for 6 hours, collected, washed with PBS-T, and samples frozen in liquid nitrogen. Frozen samples were stored at -80°C until processing.

### ChIP Protocol

ChIP-seq sample preparation protocol was modeled from *Sen et al.* with modifications (64). Frozen worms were lysed by a pre-chilled biopulverizer (Biospec, #59013N). Pulverized samples were moved to a pre-chilled mortar and grinded vigorously with mortar and pestle (Fisher, #FB961B; #FB961L). Sample powder was moved to tubes and reconstituted in Nuclei Preparation Buffer (NPB) (10 mM HEPES pH 7.5, 10 mM KCl, 1.5 mM MgCl_2_, 0.1% Triton X-100) containing protease inhibitors (Promega, #G6521). Samples were cross-linked in 1.1% PFA (Fisher, #50-980-487) for 4 minutes and quenched for 10 minutes with 125 mM (final concentration) Glycine (VWR, #97061-128). Samples were washed in NPB + protease inhibitors and resuspended in lysis buffer (50 mM Tris pH 8.1, 10 mM EDTA, 0.25% SDS) plus protease inhibitors. A Diagenode Bioruptor Plus sonicator (#B01020001) and TPX tubes (Diagenode, #NC0065146) were used to sonicate samples to generate 400-500 bp DNA fragments. Sonicated samples were centrifuged, supernatant removed, and its protein concentration determined by Bradford analysis (BioRad, #5000006). Samples were taken as a protein and DNA input control. The immunoprecipitation was performed by calculating the volume of sample needed for 1 mg of total input protein. This volume was diluted by half using broad nuclei lysis buffer (10 mM Tris-HCl pH 7.5, 1% Triton X-100, 0.5% Na Deoxycholate, 0.1% SDS). ChIP dilution buffer (16.7 mM Tris-HCl pH 8.1, 1.1% Triton X-100, and 167 mM NaCl, 1.2 mM EDTA, 0.01% SDS) was added to obtain a final IP volume of 2 mls. 2 μg of anti-V5 mouse monoclonal antibody (R&D Systems Bio-techne-NBP1-80562) was added and incubated overnight at 4°C with rotation. The next morning, Protein G dynabeads (Invitrogen, #10-003-D) were washed with blocking buffer (1x PBS+0.5% TWEEN, 0.5% BSA plus protease inhibitor) and added to IP samples and incubated for 1 hour at 4°C with rotation. IP samples were washed with RIPA low (20 mM Tris-HCl pH 8.1, 140 mM NaCl, 1 mM EDTA, 0.1% SDS, 1% Triton x-100, 0.1% DOC), RIPA high (20 mM Tris-HCl pH 8.1, 500 mM NaCl, 1 mM EDTA, 0.1% SDS, 1% Triton x-100, 0.1% DOC), LiCl (10 mM Tris-HCl pH 8.1, 250 mM LiCl, 1 mM EDTA 0.5%, Triton X-100, 0.5% DOC), and TE buffer (10 mM Tris-HCl pH 8.0, 1 mM EDTA). Samples were placed into elution buffer (10 mM Tris-HCl pH 8.0, 300 mM NaCl, 5 mM EDTA, 0.1% SDS, 5 mM DTT) containing 5 mg/mL Proteinase K (NEB, #P8107S) and 62.5 ug/mL RNAse A (Sigma, # R6513-50MG), and cross-links were reversed overnight by incubation at 37°C. DNA was purified using AMPure XP beads (Beckman Coulter, #A63880) protocol, and concentration was determined by Qubit analysis (Thermo, #Q33238). Input and IP sequencing libraries were generated using the NEBNext® Ultra™ II DNA Library Prep Kit for Illumina® (NEB, #E7645S) with barcodes unique to each of the samples (NEB, #E7335S; #E7500S). Samples were pooled and sequenced using the NextSeq 2000 (200 cycles, 400M total reads) at the Indiana University Center for Medical Genomics Core Facility. All ChIP-seq experiments were repeated at least three times.

### Sequencing Analyses

Fastq raw sequencing files were assessed for quality and filtered to remove adaptors by fastp (65). Filtered reads were aligned to the *C. elegans* genome using Bowtie 2 (66) with WBcel235(*ce11*) as the reference genome. Read alignments were further processed using SAMtools (67) to remove duplicate and mitochondrial reads and BEDtools (68) to remove any “blacklisted” alignments found in the *C. elegans* genome (UCSC). Samples with low read counts were excluded from downstream analysis: IP samples with read count of less than 15 million and input samples with read count less than 9 million. DeepTools (69) was used to generate bigwig files for visualization of enriched regions on genome browsers (**Fig 4B**). To visualize chromatin incorporation, enriched regions were identified using epic2, a program efficient at identifying diffuse ChIP-seq domains (70). Background enriched regions were identified in N2 controls and filtered from histone samples using BEDtools (68). All filtered enriched regions from histone samples were used to generate heatmaps to visualize the genome-wide enrichment profile of specific histone variants using Deeptools (**Fig 3B**) (69). Multidimensional Scaling (MDS) analysis was performed in edgeR. Enriched regions called by epic2 in all samples (*his-58*, *his-9*, and N2 IP control) were combined into a union set using BEDtools and reads in all enriched regions counted by featureCounts (71). Count files were then used in edgeR to generate MDS plot (**Fig 3C**). Genomic targets of histone variants in fed and starved samples were identified by performing edgeR differential analysis comparing reads from histone reporter samples to inputs and N2 IgG controls (72), applying a cutoff of a log_2_ fold change (log_2_FC) greater than 1 and false discovery rate (FDR) of less than or equal to 0.05. Read counts at each protein coding gene locus were determined using featureCounts with the protein coding gene coordinates as the input (73, 74). Tissue enrichment (TEA) and Gene Ontology (GO) analyses was performed using the Wormbase Bioinformatics tool (**Fig 4C, D**) (75). For differential analysis between fed and starved histone reporters, the genomic coordinates of all *C. elegans* protein coding genes was used as an identifier (73, 74). The number of reads on all protein coding genes in each sample was determined using featureCounts (71). Reads between fed and starved samples were compared using edgeR (72), applying a cutoff of a log_2_ fold change (log_2_FC) greater than 1 and false discovery rate (FDR) of less than or equal to 0.05 to identify gene targets with significant enrichment changes.

## Acknowledgements

The authors thank D. Rusch of the Center for Genomics and Bioinformatics (Indiana University Bloomington) for help with ChIP-seq analyses and members of the Aoki and Roh laboratories for helpful discussion. The Center for Medical Genomics at Indiana University School of Medicine assisted in DNA sequencing and is partially supported by the Indiana University Grand Challenges Precision Health Initiative (PHI). H.R. is supported by the National Institute of Diabetes and Digestive and Kidney Diseases (R01DK129289) and American Diabetes Association Junior Faculty Award (7-21-JDF-056). S.T.A. and this work was funded by the PHI, Showalter Trust, and NIH/NIGMS (R35GM142691).

## Supplemental Figure Captions

**Figure S1.**
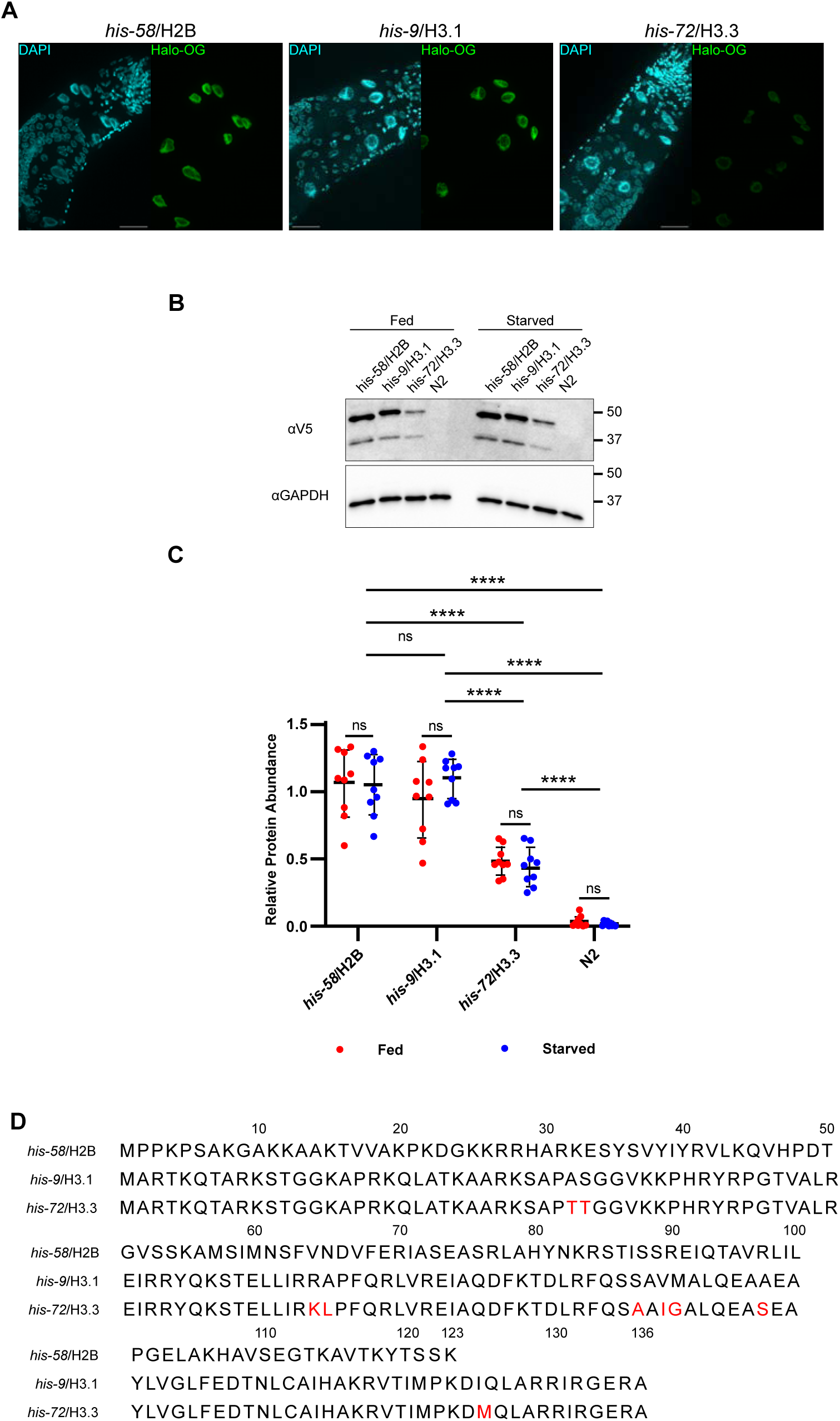
Additional histone reporter data. (**A**) Representative images of anterior portion of histone reporter worm intestines stained with DAPI and Halo-Oregon Green (Halo-OG). Images shown as max intensity projection of confocal stack taken at 63X magnification. Scale bar, 25 µM. (**B**) Immunoblot comparing histone reporter expression between fed and starved strains. Synchronized, young adult histone reporter worms were either fed or starved for 6 hours prior to sample collection. 110 worms were collected for each sample (n=3). 3 dilutions of protein lysate were loaded for each sample. (**C**) Quantification of histone reporter protein expression between fed and starved strains. Quantification performed using ImageJ. Area of histone reporter band normalized to GAPDH loading control. Statistical significance was determined using a two-way ANOVA with Tukey’s Multiple Comparison Test. ****p<0.0001; ns, not significant. (**D**) Amino acid sequences of the histone proteins used in this study. Amino acids highlighted in red of *his-72*/H3.3 reporter indicate sequence differences compared to *his-9*/H3.1.

**Figure S2.**
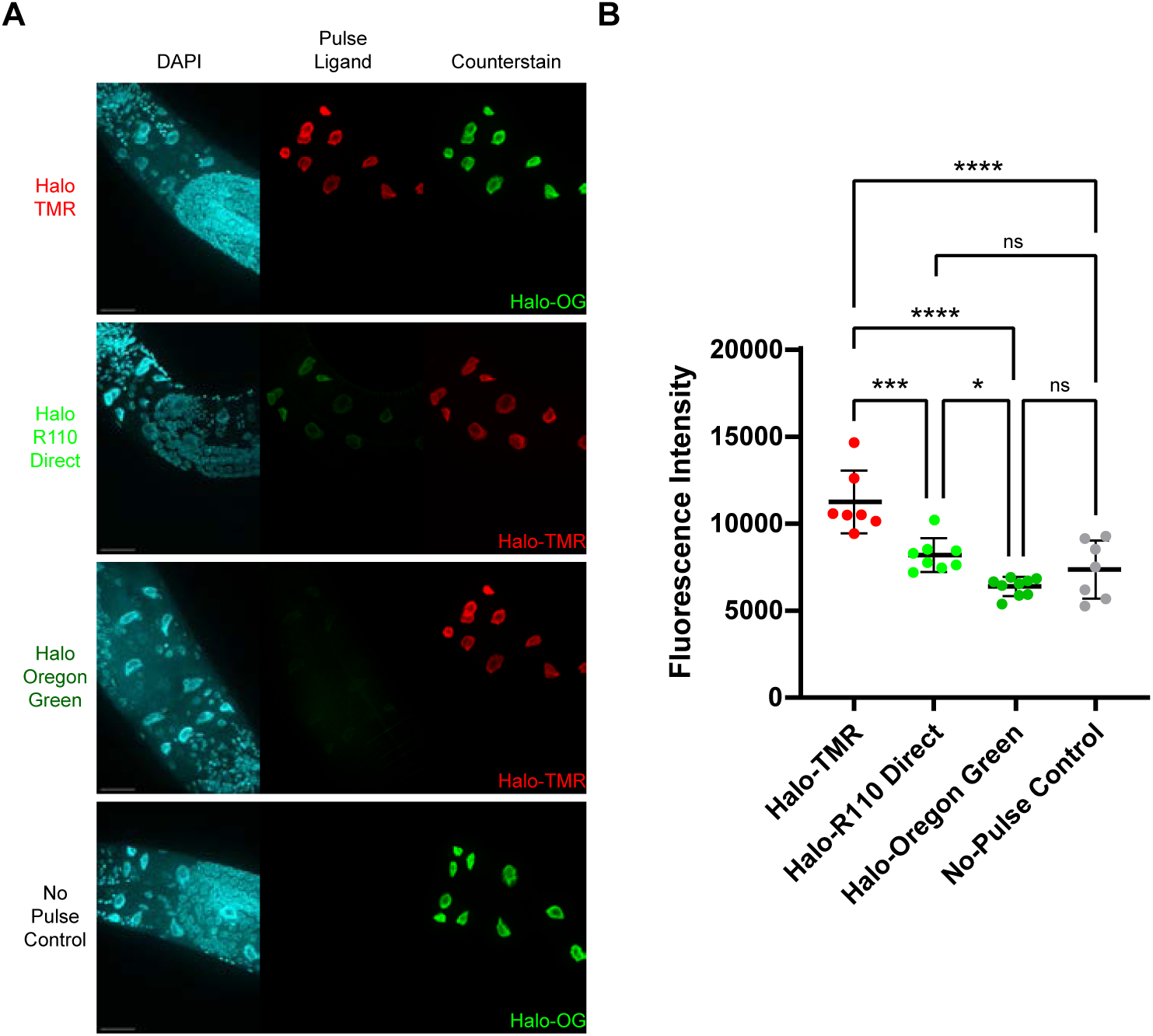
In vivo labeling of an intestinal histone reporter with commercially available HaloTag ligands. (**A**) Representative images of *his-58*/H2B reporter worms pulse-labeled with either Halo-TMR, Halo-R110Direct, or Halo-Oregon Green. See **Results** and **Methods** for more details. Images shown as max intensity projection of confocal stack taken at 63X magnification under oil immersion. Scale bar, 25 µm. (**B**) Fluorescence quantitation of *his-58*/H2B worms pulse-labeled with Halo ligands. Significance was determined by performing a two-way ANOVA with Tukey’s multiple comparisons test. *p<0.05; ***p<0.001; ****p<0.0001; ns, not significant.

**Figure S3.**
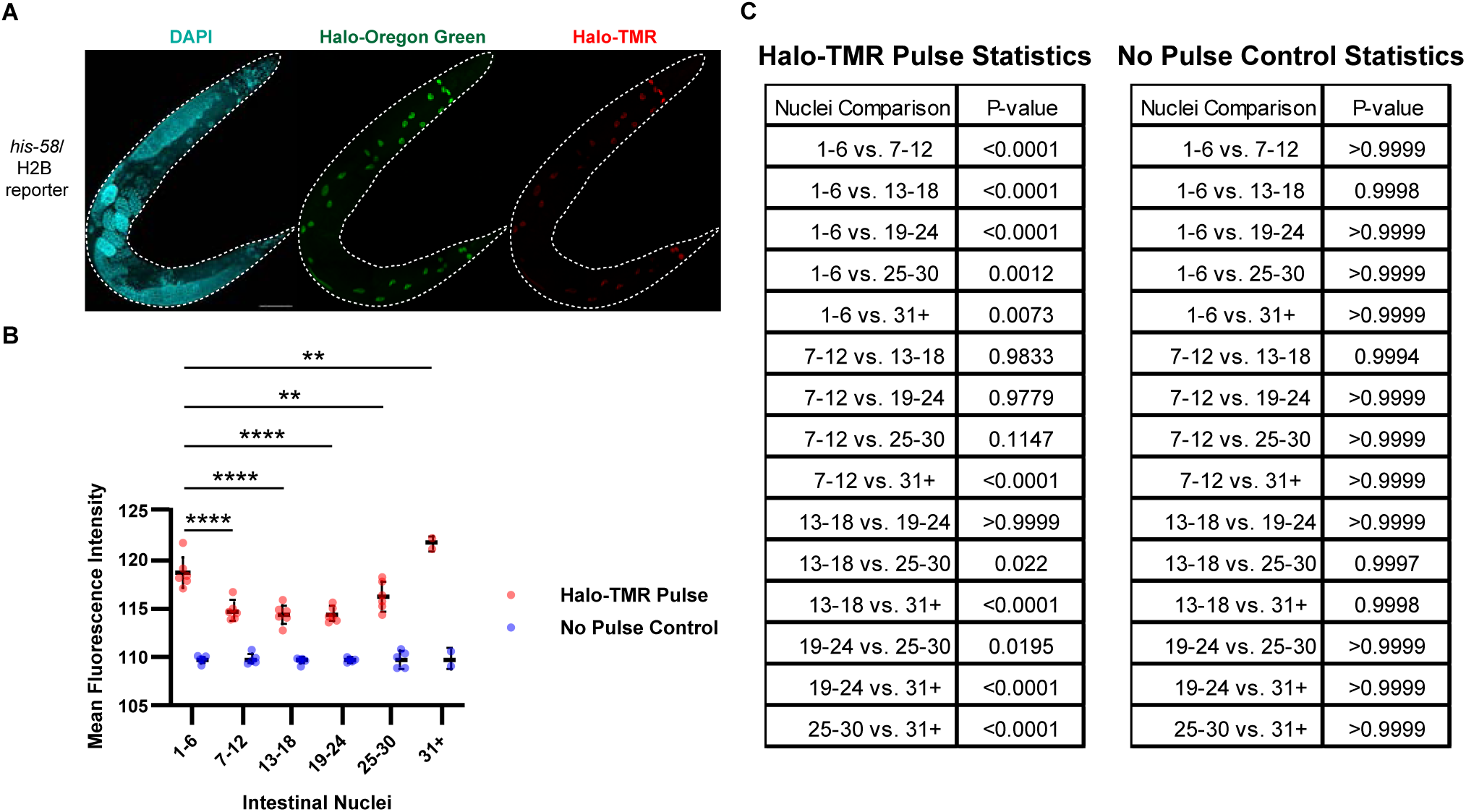
Anterior nuclei are labeled most efficiently by soaking. (**A**) Representative images of *his-58*/H2B reporter worms pulse-labeled with 300 nM Halo-TMR and immediately fixed, stained with DAPI and Halo-Oregon Green, and imaged. Images shown as montage of max intensity projection of confocal stack taken at 40x magnification under oil immersion. Scale bar, 75 µm. (**B**) Quantitation of Halo-TMR fluorescence. Significance was determined by performing a two-way ANOVA with Tukey’s multiple comparisons test. **p<0.01, ****p<0.0001. (**C**) Summary statistics from two-way ANOVA with Tukey’s multiple comparisons test on nuclei groupings from Halo-TMR pulse and No pulse control samples.

**Figure S4.**
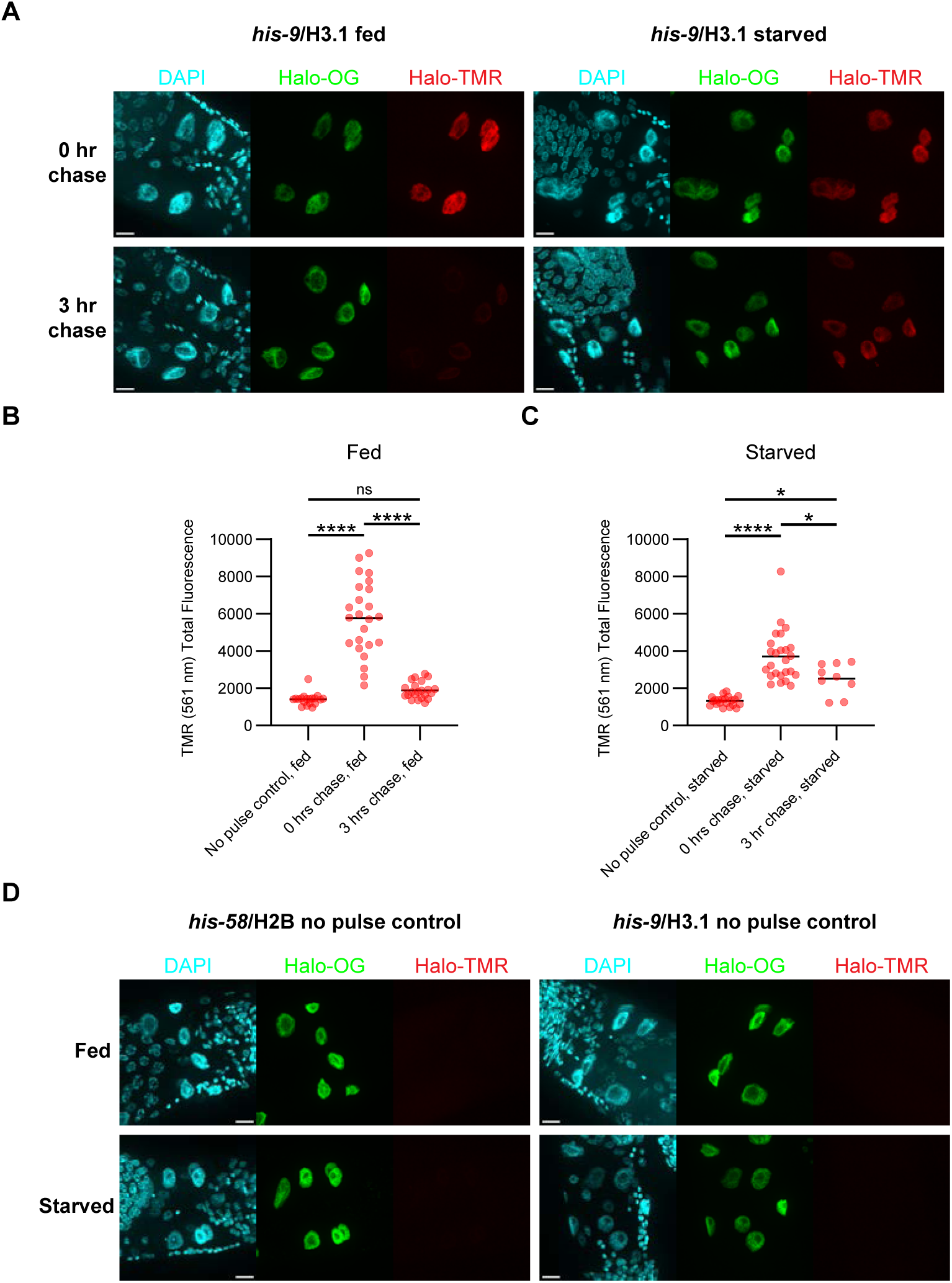
In vivo pulse-chase labeling of intestinal *his-9*/H3.1 histone reporter worms. (**A**) Representative images of fed and starved *his-9*/H3.1 reporter anterior intestinal nuclei after a 0- or 3-hour chase. Images shown as max intensity projection of confocal stack taken at 63X magnification under oil immersion. Scale bar, 10 µm. (**B-C**) Fluorescence intensity measurements of fed (B) and starved (C) *his-9*/H3.1 reporter anterior nuclei (see **Results** and **Methods**). Significance was determined by performing a two-way ANOVA with Tukey’s multiple comparisons test. *p<0.05; ****p<0.0001; ns, not significant. (**D**) Representative images of fed and starved *his-58*/H2B and *his-9*/H3.1 no pulse control samples. Images shown as max intensity projection of confocal microscopy stack taken at 63X magnification. Scale bar, 10 µm.

**Figure S5.**
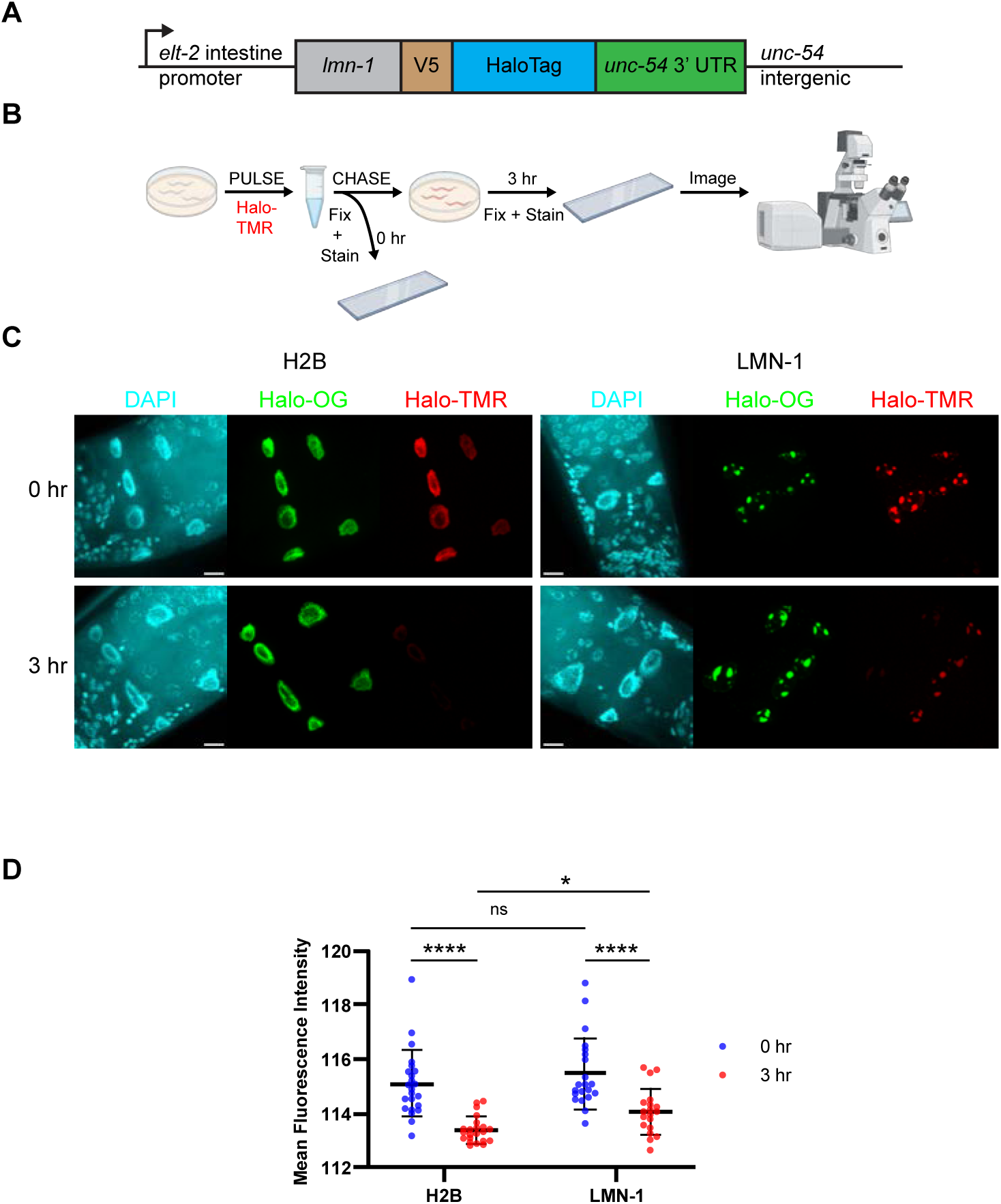
LMN-1 reporter is more stable than H2B reporter using Halo tag *in vivo* pulse-chase method. (**A**) Linear diagram of the HaloTag LMN-1 reporter. An *elt-2* intestine-specific promoter drives expression of LMN-1 fused to a V5 epitope and HaloTag. The reporter has an *unc-54* 3’UTR and intergenic region. (**B**) HaloTag pulse-chase method. Reporter worms were pulse-labeled by soaking reporter worms in Halo-TMR ligand. Worms were either collected immediately (0 hr) or moved back to plates for 3 hours (3 hr). At each timepoint, worms were fixed and stained to visualize nuclei and total reporter protein. Worms were imaged on a confocal microscope and Halo-TMR fluorescence used as a measure of protein stability. (**C**) Representative fluorescent images of reporter worms collected at each timepoint stained with DAPI, Halo-Oregon Green (Halo-OG), and Halo-TMR. Scale bar, 10 µm. (**D**) Quantification and analysis of Halo-TMR fluorescence intensity in reporter worms. Significance was determined by performing a two-way ANOVA with Tukey’s multiple comparisons test. *p<0.05; ****p<0.0001; ns, not significant.

**Figure S6.**
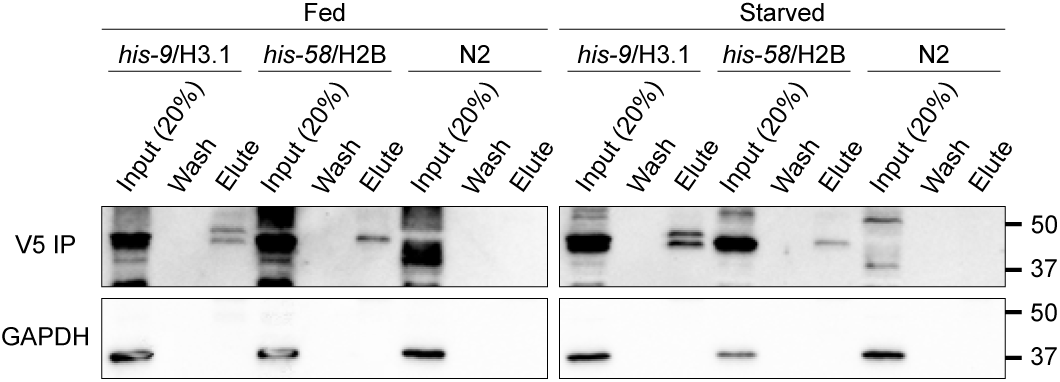
Chromatin Immunoprecipitation analysis. Immunoblot of chromatin immunoprecipitation (ChIP) of histone reporter proteins. Input samples loaded as a 20% ratio of elute sample to evaluate efficiency of immunoprecipitation. Histone reporter proteins were successfully immunoprecipitated and enriched.

**Table S1.**
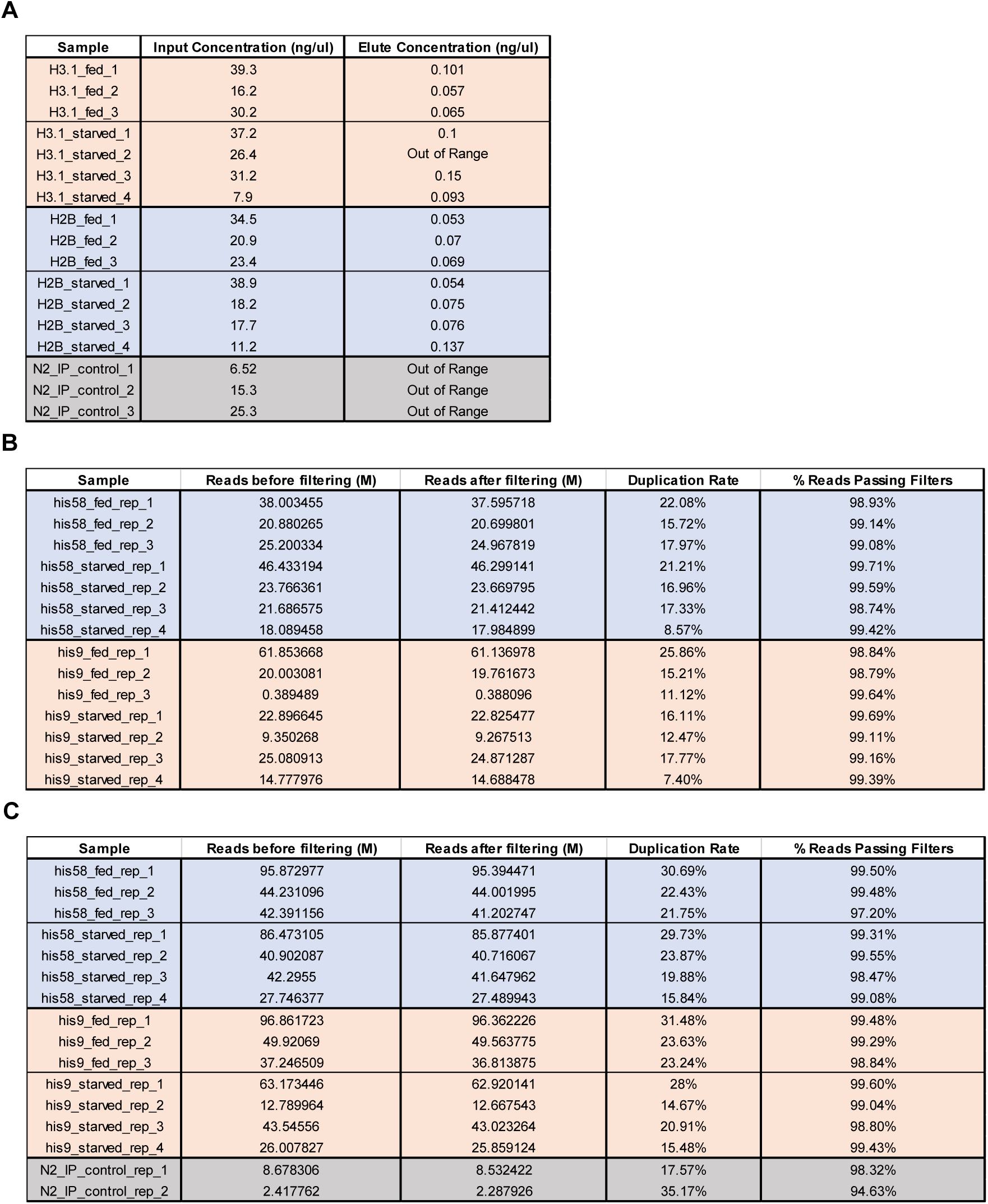
ChIP-seq DNA metrics. (**A**) Quantification of DNA concentration in ChIP-seq experimental samples. DNA concentrations were comparable in all inputs. In IP samples, DNA was detected in histone samples but undetectable in N2 IP control samples. DNA quantification performed with a Qubit dsDNA high sensitivity kit. (**B**) Raw read metrics from input samples filtered with Fastp and featureCounts programs. (**C**) Raw read metrics from IP samples filtered with Fastp and featureCounts programs.

| Sample | Input Concentration (ng/ul) | Elute Concentration (ng/ul) |
| --- | --- | --- |
| H3.1_fed_1 | 39.3 | 0.101 |
| H3.1_fed_2 | 16.2 | 0.057 |
| H3.1_fed_3 | 30.2 | 0.065 |
| H3.1_starved_1 | 37.2 | 0.1 |
| H3.1_starved_2 | 26.4 | Out of Range |
| H3.1_starved_3 | 31.2 | 0.15 |
| H3.1_starved_4 | 7.9 | 0.093 |
| H2B_fed_1 | 34.5 | 0.053 |
| H2B_fed_2 | 20.9 | 0.07 |
| H2B_fed_3 | 23.4 | 0.069 |
| H2B_starved_1 | 38.9 | 0.054 |
| H2B_starved_2 | 18.2 | 0.075 |
| H2B_starved_3 | 17.7 | 0.076 |
| H2B_starved_4 | 11.2 | 0.137 |
| N2_IP_control_1 | 6.52 | Out of Range |
| N2_IP_control_2 | 15.3 | Out of Range |
| N2_IP_control_3 | 25.3 | Out of Range |

**B**
| Sample | Reads before filtering (M) | Reads after filtering (M) | Duplication Rate | % Reads Passing Filters |
| --- | --- | --- | --- | --- |
| his58_fed_rep_1 | 38.003455 | 37.595718 | 22.08% | 98.93% |
| his58_fed_rep_2 | 20.880265 | 20.699801 | 15.72% | 99.14% |
| his58_fed_rep_3 | 25.200334 | 24.967819 | 17.97% | 99.08% |
| his58_starved_rep_1 | 46.433194 | 46.299141 | 21.21% | 99.71% |
| his58_starved_rep_2 | 23.766361 | 23.669795 | 16.96% | 99.59% |
| his58_starved_rep_3 | 21.686575 | 21.412442 | 17.33% | 98.74% |
| his58_starved_rep_4 | 18.089458 | 17.984899 | 8.57% | 99.42% |
| his9_fed_rep_1 | 61.853668 | 61.136978 | 25.86% | 98.84% |
| his9_fed_rep_2 | 20.003081 | 19.761673 | 15.21% | 98.79% |
| his9_fed_rep_3 | 0.389489 | 0.388096 | 11.12% | 99.64% |
| his9_starved_rep_1 | 22.896645 | 22.825477 | 16.11% | 99.69% |
| his9_starved_rep_2 | 9.350268 | 9.267513 | 12.47% | 99.11% |
| his9_starved_rep_3 | 25.080913 | 24.871287 | 17.77% | 99.16% |
| his9_starved_rep_4 | 14.777976 | 14.688478 | 7.40% | 99.39% |

**C**
| Sample | Reads before filtering (M) | Reads after filtering (M) | Duplication Rate | % Reads Passing Filters |
| --- | --- | --- | --- | --- |
| his58_fed_rep_1 | 95.872977 | 95.394471 | 30.69% | 99.50% |
| his58_fed_rep_2 | 44.231096 | 44.001995 | 22.43% | 99.48% |
| his58_fed_rep_3 | 42.391156 | 41.202747 | 21.75% | 97.20% |
| his58_starved_rep_1 | 86.473105 | 85.877401 | 29.73% | 99.31% |
| his58_starved_rep_2 | 40.902087 | 40.716067 | 23.87% | 99.55% |
| his58_starved_rep_3 | 42.2955 | 41.647962 | 19.88% | 98.47% |
| his58_starved_rep_4 | 27.746377 | 27.489943 | 15.84% | 99.08% |
| his9_fed_rep_1 | 96.861723 | 96.362226 | 31.48% | 99.48% |
| his9_fed_rep_2 | 49.92069 | 49.563775 | 23.63% | 99.29% |
| his9_fed_rep_3 | 37.246509 | 36.813875 | 23.24% | 98.84% |
| his9_starved_rep_1 | 63.173446 | 62.920141 | 28% | 99.60% |
| his9_starved_rep_2 | 12.789964 | 12.667543 | 14.67% | 99.04% |
| his9_starved_rep_3 | 43.54556 | 43.023264 | 20.91% | 98.80% |
| his9_starved_rep_4 | 26.007827 | 25.859124 | 15.48% | 99.43% |
| N2_IP_control_rep_1 | 8.678306 | 8.532422 | 17.57% | 98.32% |
| N2_IP_control_rep_2 | 2.417762 | 2.287926 | 35.17% | 94.63% |

**Table S2.**
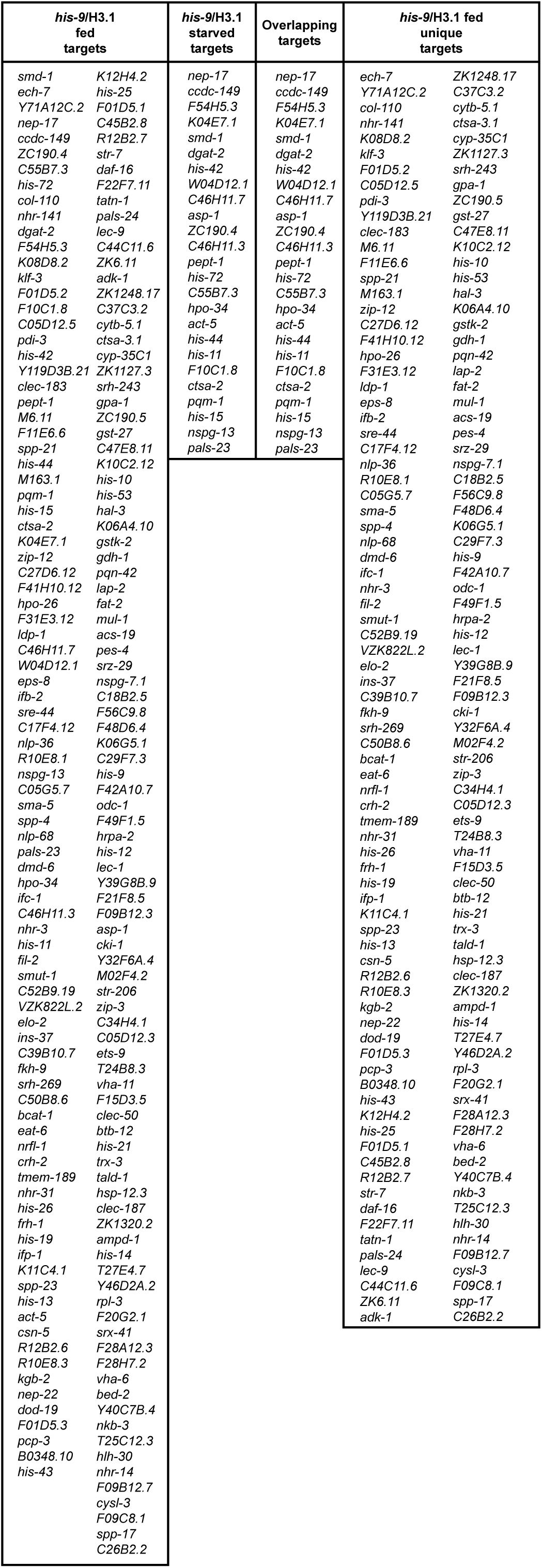
*his-9*/H3.1 reporter gene targets identified by ChIP-seq. 187 genes were identified as *his-9*/H3.1 targets in fed animals and 25 genes were identified as *his-9*/H3.1 targets in starved animals. All 25 starved targets were found in fed samples.

**Table S3.** Strains and plasmids used in this study.

**A**
| Strain | Genotype | Description |
| --- | --- | --- |
| N2 Bristol |  | Wildtype strain |
| EG6699 | *ttTi5605 II; unc-119(ed3) III; oxEx1578* | MosSCI worm generation |
| CDE5 | *ddcSi1 [elt-2 promoter::his-9::linker::1xV5::Halo opt with introns::unc-54 3'UTR and intergenic] II* | *his-9* reporter strain |
| CDE6 | *ddcSi2 [elt-2 promoter::his-58::histone linker::1xV5::Halo opt with introns::unc-54 3'UTR and intergenic] II* | *his-58* reporter strain |
| CDE8 | *ddcSi4 [elt-2 promoter::his-72::histone linker::1xV5::Halo opt with introns::unc-54 3'UTR and intergenic] II* | *his-72* reporter strain |
| CDE142 | *ddcSi43 [elt-2 promoter::lmn-1::histone linker::1xV5::Halo opt with introns::unc-54 3'UTR and intergenic] II* | *lmn-1* reporter strain |

**B**
| Plasmid | Description |
| --- | --- |
| pCFJ151 | Standard Multiple Cloning site vector to generate MosSCI inserts at ttTi5605 location. |
| pGH8 | Coinjection marker. mCherry expression in the nervous system. |
| pCFJ90 | Use as co-injection marker. Red mCherry fluorescence in the pharynx. |
| pCFJ104 | Red mCherry expression in the pharynx driven by the myo-3 promoter. |
| pCFJ601 | Mos1 Transposase |
| pDDC14 | MosSCI vector, *elt-2* promoter, *his-58* tagged with V5;Halo, *unc-54* 3'UTR and intergenic |
| pDDC15 | MosSCI vector, *elt-2* promoter, *his-9* tagged with V5;Halo, *unc-54* 3'UTR and intergenic |
| pDDC16 | MosSCI vector, *elt-2* promoter, *his-72* tagged with V5;Halo, *unc-54* 3'UTR and intergenic |
| pDDC171 | MosSCI vector, *elt-2* promoter, *lmn-1* tagged with V5;Halo, *unc-54* 3'UTR and intergenic |

